# Estimating the cortex-wide overlap between wordless narrative scene comprehension, reading comprehension, and topological visual, auditory, and somatomotor maps

**DOI:** 10.1101/264002

**Authors:** Mariam R. Sood, Martin I. Sereno

## Abstract

Previous imaging research in scene comprehension has mostly focussed on isolated scenes and objects. Here we use narrative picture stimuli and a novel controlled presentation technique to identify the full extent of activation during naturalistic narrative scene comprehension. We then situate it with respect to topologically-mapped sensory and motor regions as well as cortical regions activated during naturalistic narrative reading comprehension. The data for all experiments was obtained from the same set of subjects, and analyzed using cortical surface based methods from start to finish. The results suggest that scene and reading activations are spread across occipital, parietal, temporal, and frontal cortex, and largely aligned with each other. Within these regions, there were also sites uniquely activated when subjects were engaged in either narrative scene comprehension or in reading comprehension. Finally, the cortical activations for both scene and reading comprehension contrasts substantially overlap topological cortical maps, specifically retinotopic and tonotopic maps in occipital, parietal, temporal, and frontal cortex.

---

What is the neural basis of sequential non-verbal visual comprehension? We know that humans generate and process the results of long sequences of fixations in rich visual environments from the time of their birth. By contrast, a majority of the research in scene comprehension has focussed on isolated objects and pictures. So far several scene selective regions have been identified in the occipital and inferior temporal cortex. Among them, the two best known are the parahippocampal place area (PPA) in the collateral sulcus near the parahippocampal-lingual boundary (Epstein & Kanwisher, 1998) and the retrosplenial complex (RSC) (Bar & Aminoff, 2003). A third region, the occipital place area (OPA) (Dilks et al., 2013) has been found near the transverse occipital sulcus. All three of these regions respond preferentially to pictures depicting scenes, spaces, and landmarks compared to pictures of faces or single movable objects (for a review, see Epstein & MacEvoy, 2011). Similarly, the fusiform face area, a region in the mid fusiform gyrus (Sergent et al., 1992; Kanwisher et al., 1997) and the occipital face area, a region in lateral occipital cortex in the vicinity of inferior occipital gyrus (Puce et al, 1996; Yovel and Kanwisher, 2005) show preferential selectivity to faces, while a region in lateral occipital cortex inferior to V3A has been shown to prefer objects over scrambled objects.

Only a few studies have looked across the whole brain while participants watched more naturalistic sequential visual stimuli such as movies (Bartels and Zeki, 2004; Hasson et al., 2004, 2008; Nishimoto et al., 2011, Huth et al., 2012). These studies have shown that the BOLD activity evoked by natural movies extends well beyond occipital cortex and is spread across temporal, parietal, and frontal cortex. While there has been some effort to analyze the retinotopic map architecture underlying scene selective regions in the occipital cortex, no studies so far have looked at scene selective regions in relation to topological sensory-motor maps across all higher level cortices, using the same subjects.

In the study presented here, the same set of subjects took part in a narrative scene comprehension experiment, a narrative reading comprehension experiment, and then topological visual, auditory and somatomotor mapping experiments. The narrative scene comprehension experiment consisted of a ‘picture based narrative comprehension’ fMRI task. Subjects viewed a series of pictures adapted from wordless picture story books. To precisely control fixation location and duration across all conditions, participants' saccades across each single- or multi-frame picture page were directed by ‘saccading’ a transparent gaussian ‘bubble’-style mask (at 1 Hz) to relevant points in the images chosen by offline comprehension testing. The narrative reading comprehension experiment used an analogous technique to control and direct saccades by revealing only one word at a time in sequence (other words replaced by gray rectangles) with each word in its natural reading position. The reading comprehension experiment and the sensory and motor mapping studies are described in more detail in Sood & Sereno (2016). Thus in both the narrative scene comprehension experiment and the reading comprehension experiment, participants were required to comprehend a narrative sequence (using pure picture scenes or words), and eye movements within each experiment were matched and controlled across all experimental conditions.

There were two primary objectives behind this study. Firstly, to site narrative scene comprehension regions relative to topological retinotopic, tonotopic and somatomotor maps across the whole cortex. While initially it was thought that the scene/object selective regions in occipital cortex fell beyond the bounds of retinotopy, subsequent discoveries in retinotopic maps have shown that most of the scene/object selective regions in the posterior occipital cortex fall within or adjacent to retinotopic maps. There has not been a systematic effort to localize the higher level (beyond occipital cortex) scene processing regions relative to topological sensory-motor maps discovered in frontal, parietal and temporal lobe. This study sheds light on the underlying neural organization of these regions and allows us to accurately site these regions with respect to topological map borders that can be independently demarcated.

Secondly, by combining this data with the reading comprehension data (and the topological sensory-motor maps), we can draw stronger inferences regarding the degree to which the serial assembly processes in linguistic and non-linguistic comprehension using the same modality intersect and diverge. Scene comprehension, like language, is fundamentally serial in nature. The integration of successive glances in the comprehension of a visual scene (and even more in a series of pictures) requires a kind of serial assembly operation similar to the serial integration of word meaning in language comprehension. An isolated glance taken out of scene context is as ambiguous as a single word taken out of discourse context (Sereno, 2014). Although a linguistic task such as reading and a non-linguistic task such as narrative scene comprehension involve cognitive processes unique to each (orthographic, phonological, lexical-syntactic for reading; object processing, relation-to-background processing, gist processing in scene comprehension), there are also several processes that could very well be shared between the two, such as semantic access, event segmentation, discourse structure building, and so on. We hypothesized that some of the activations identified in reading comprehension (Sood & Sereno, 2016) -- especially in frontal cortex -- might be shared with the activation observed during this non-linguistic visual task. We were also interested whether we could identify frontal and other higher level regions often thought to be domain general that were in fact unique to either scene comprehension or reading.

On the methodological front, all five data sets (Scene Comprehension, Reading, Visual, Auditory, and Somatomotor maps) were processed with a start-to-finish surface-based group analysis, which has been shown to provide better spatial resolution as a direct result of avoiding blurring across sulci (Fischl et al., 1999a,b). This is particularly relevant when attempting to accurately measure overlap.

## Materials and Methods

### Subjects

The data presented here comes from 20 right-handed native English speakers (9 women). The mean age was 28 (ranging from 19 to 58). All participants were neurologically healthy with normal or corrected to normal vision and normal hearing capacity. The experimental protocols were approved by local ethics committees and participants gave their informed written consent prior to the scanning session. The study required each participant to take part in five separate fMRI experimental sessions: narrative scene comprehension, reading, retinotopic mapping, auditory mapping and somatomotor mapping. The experimental design and results from four of these experiments -- reading, and Retinotopic, Auditory and Somatomotor mapping -- were detailed in a previous publication (Sood & Sereno, 2016). All 20 participants took part in the Reading task and Retinotopic mapping experiments. 18 of the same participants took part in auditory mapping and 17 of the same participants took part in Somatomotor mapping. From the same set of subjects, 9 participated in the narrative Scene Comprehension task.

### Experimental Stimuli and design

#### Narrative Scene comprehension Experiment

The picture based comprehension experiment (Figure 1) consisted of comprehending a coherent story from a series of pictures (no text captions, 'bubbles', or stray text of any kind). The stories presented in the experiment came from 12 wordless picture story books. Any incidental text in the pictures was edited out in Photoshop. Each page of the book was not presented in its entirety. Instead, a transparent gaussian 'bubble'-style mask (fwhm: 512 pixels) was moved in a saccadic fashion over relevant parts of the page (1920 by 1080 pixels) at 1 Hz. The subjects were instructed to move their eyes along with the mask center to follow the story on the page. The mask locations were carefully chosen by offline comprehension testing. Each book's presentation time was thus determined by the total number of mask/saccade locations across all pages. The book presentation times varied between 30-50 seconds. Each book was presented in a separate run (12 runs in total). In each run, 3 conditions and a central fixation screen were presented.

**Figure 1:**
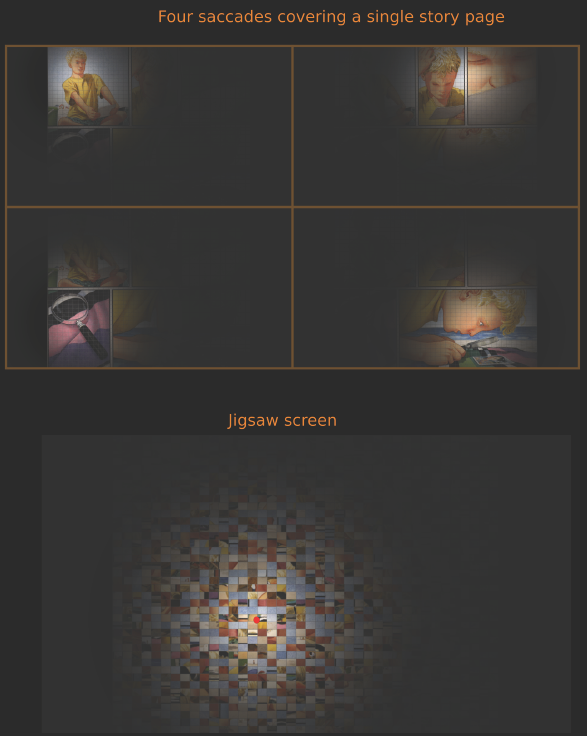
Stimulus screens.

In the main experimental condition (Story), the pictures were presented in the same order as they appeared in the books. In the second condition (Jigsawed), a jigsawed version (each page fragmented into 40×40 pixel rectangular blocks and the blocks shuffled) of each book page was presented. Although the page order remained the same, the jigsaw made the book pages completely incomprehensible. The same Gaussian ‘bubble’ mask locations as those used for the Story condition were used. In the third condition (Shuffled), the presentation order of the scenes in the book was shuffled. In the Shuffled condition, the subjects eventually saw all the scenes that were seen in the Story condition, but the temporally coherent story structure was disrupted by shuffling. Since the jigsawed images have a rectangular grid pattern, which results in high spatial frequency content, a 40×40 pixel grid pattern made of orthogonal gray lines was superimposed on all stimuli. The length of each of the three conditions, and the number of mask locations was the same for each book. Before the start of each condition, a ready screen was presented for 2 seconds. The Story and the Shuffled conditions were always separated by a Jigsawed condition. In roughly half the runs (selected randomly) the Story condition came before Jigsawed while in the rest, the Shuffled condition came before Jigsawed. The central fixation screen (15 seconds in duration) was presented at a random block position within the run. For all three experimental conditions, participants were instructed to saccade along with the mask center location.

The stimulus presentation mode used here, where different parts of the picture page was made visible briefly, served several purposes. In the spirit of the classic attention study by Posner (1980), the subject’s exogenous attention is automatically drawn towards the next highlighted, higher-contrast region in the peripheral visual field, thus ensuring exogenous attentional control across all experimental conditions. This presentation mode also ensured that participants made controlled eye saccades -- and extremely similar eye movements across conditions -- as opposed to the variable, uncontrolled eye movements that would occur if the picture pages were presented unmasked for free viewing. Additionally, this presentation mode ensured that different subjects had much more similar viewing experiences when comprehending the picture stories.

In order to further ensure that participants stayed attentive and made similar eye movements for all conditions, they were asked to press a button when a red dot was presented (at random) at the centre of the gaussian mask. When the task occurred in the fixation screen, the fixation dot turned red for a duration of 1 second. The level of comprehension achieved for the stories were measured with a questionnaire afterwards. Participants were informed of this quality control process before the scan.

The stimulus presentation framework was programmed in C/OpenGL/X11 (Mac/Linux binaries available on request). The stories were adapted from the following 12 wordless picture books: Flotsam by David Wiesner, Journey by Aaron Becker, Freefall by David Wiesner, Tuesday by David Wiesner, Goodnight Gorilla by Peggy Rathmann, The Grey Lady and the Strawberry Snatcher by Molly Bang, Window by Jeannie Baker, Oops by Arthur Geisert, Belonging by Jeannie Baker, Changes Changes by Pat Hutchins, Deep in the Forest by Brinton Turkle and Pancakes for Breakfast by Tomie dePaola.

Although this task involved a lot more than the comprehension of an isolated scene, for simplicity of description, we will use ‘Scene Comprehension’ to describe the task in the rest of this paper.

#### Narrative reading comprehension Experiment

The reading experiment (fully described in Sood & Sereno, 2016) used short narrative comprehension passages in English (64 words, ~4 words/sec, word duration a function of word length) shown one word at a time with each word in its natural reading position. Other words were presented as grayed rectangles. Contrast conditions were meaningless (to the subjects, or in Hindi) same-character-length Hindi character strings, or a large dot, and finally, a central fixation screen as OFF. The experiment consisted of four runs, where each run comprised 32 blocks presented in a random order. To control for variations in attention across conditions, a secondary task was to press a button when the color of an English word, Hindi 'word', large dot, or fixation changed from black to grey. The level of comprehension achieved for each unrelated English passage was measured with a questionnaire afterward.

#### Cortical mapping experiments

The cortical mapping experiments carried out were detailed in Sood & Sereno, 2016. Retinotopic mapping and auditory mapping experiments closely followed previous work (Sereno et al., 2013, Dick et al., 2012). In somatomotor mapping, participants moved 11 body parts progressing from tongue to toe following a short auditory cue. Runs (eight 64s cycles; 4 runs in total) alternated between movement cycles in each direction (tongue to toe, toe to tongue).

### Experimental set-up

The stimuli were back projected at HDMI (1920 x 1080 pixels) resolution onto a screen inside the bore of the magnet almost flush with the back of the head coil that was visible to the subjects via a mirror; viewing distance was 30 cm. Memory foam cushions (NoMoCo Inc.) were packed around the head to provide additional passive scanner acoustical noise attenuation and to stabilize head position. Responses were made via an optical-to-USB response box (LUMItouch, Photon Control, Burnaby, Canada) situated under their right hand. We used a 30-channel head coil, with the eye coils removed and RC terminated (as opposed to the standard 32-channel head coil) for these scans. The lack of eye coils greatly improved the viewing experience (preventing disruptive saccade-direction-dependent blocking of the view of one or the other eye) without affecting the signal-to-noise in any part of the brain except for a slight reduction at the extreme tip (< 5mm) of the orbitofrontal pole.

### Imaging Parameters

Functional images were acquired on a 1.5 T whole-body TIM Avanto System (Siemens Healthcare), at the Birkbeck / University College London Centre for NeuroImaging (BUCNI), with RF body transmit and a 30-channel receive head coil. For all subjects, images were acquired using multiband EPI (40 slices, 3.2×3.2×3.2mm, flip=75°, TE=54.8ms, TR=1sec, accel=4) (Moeller et al., 2011). To allow longitudinal relaxation to reach equilibrium, 8 initial volumes were discarded from each run for multiband EPI. For each imaging session, a short (3 min) T1-weighted 3D MPRAGE (88 partitions, voxel resolution 1×1×2mm, flip angle=7°, TE=4ms, TI=1000ms, TR=1370ms, mSENSE acceleration=2x, slab-selective excitation) was acquired with the same orientation and slice block centre as the functional data (‘alignment scan’), for initial alignment with the high-resolution scans (acquired as part of the previous experiments) used to reconstruct the subject’s cortical surface.

### Data Analysis

The analysis of picture-story comprehension data utilized the FSL-Freesurfer cortical surface based pipeline previously described for the analysis of reading experiment data (Sood & Sereno, 2016). The overlap analysis with topological visual, auditory and somatomotor maps also followed the methods described in Sood & Sereno 2016.

#### Anatomical image processing

For each subject, the cortical surface was reconstructed with FreeSurfer (version 5; Dale et al., 1999) from the aligned average of the two high-resolution T1-weighted MPRAGE scans. Both mapping data and reading data employ a complex-valued cross-subject surface-based analysis stream that begins by sampling responses and statistics to individual reconstructed cortices (cross-subject 3D averaging was not used at any point in the pipeline).

#### Analysis of picture-story data

The single subject fMRI data was motion corrected and skull stripped using FSL tools (MCFLIRT and BET). First level fMRI analysis was carried out by applying the General Linear Model (GLM) within FEAT using FILM prewhitening (FSL, version 5) with motion outliers (detected by fsl_motion_outliers) being added as confound regressors if there was more than 1 mm motion (as identified by MCFLIRT). All subjects had minimal motion (maximum displacement well under 1 mm). The high-pass filter cut-off was estimated using the FSL Feat tool based on the power spectra of the design matrices. Three main explanatory variables were modeled and controlled: Story, Jigsawed and Shuffled. Button press responses to target 'red dot' were modeled as the fourth regressor. In order to capture slight deviations from the model, temporal derivatives of all explanatory variables convolved with FEAT's double gamma hemodynamic response function (HRF) were included. The registration from functional to anatomical (6 DOF) and standard space (12 DOF) was first done using FSL’s FLIRT and further optimized using boundary based registration (bbregister; FreeSurfer) similar to the procedure for the naturalistic reading experiment. A fixed effects analysis was performed across 12 runs from an individual subject to get group FEAT (GFEAT) results of first-level contrast of parameter estimates (COPEs) and their variance estimates (VARCOPEs) in the standard space. subject-Across group analysis was then carried out on the cortical surface using FreeSurfer tools. The GFEAT results of each subject were first sampled to individual cortical surfaces and then resampled to the spherical common average reconstructed surface (fsaverage). Surface-based spatial smoothing of 3mm FWHM was applied on the icosahedral sphere. A mixed effects GLM group analysis was performed on the average surface using the mri_glmfit program from FreeSurfer. Significance maps were thresholded at p<0.01 and were then corrected for multiple comparisons with cluster-based correction using csurf programs surfclust and randsurfclust, with clusters greater than 40 mm^2^ (on a smoothwm surface) excluded yielding a corrected significance of p<0.05. Finally corrected significance values (p<0.05) of Scene Comprehension activation were displayed on the fsaverage surface.

The single subject raw data was not spatially smoothed in 3D. For final illustrations, 5 steps (~2.2 mm FWHM) of surface-based smoothing was applied. Hence the 3D, Gaussian random field based cluster correction provided by FSL is not appropriate for multiple comparison correction of the language data. We have instead used the surface based cluster correction using surfclust/randsurfclust (Hagler et al., 2006, 2007). The GFEAT results were sampled to their respective anatomical surfaces, thresholded at p<0.001 (Z=3.09) and corrected for multiple comparisons with cortex surface clusters smaller than 30 mm2 excluded, achieving a corrected p-value of 0.01.

The target (detecting the occasional ‘red dot’) presentation timings and the button press events were logged during the experiment and analyzed to assess performance on the task. For each participant, the number of targets detected and the mean response time in each condition were calculated. The cross-subject mean response time and target detection rates were assessed for significant differences across conditions.

#### Overlap analysis

All overlaps were calculated using "original vertex-wise area" in csurf FreeSurfer. Original vertex-wise area in FreeSurfer is defined as the sum of 1/3 the area of each adjacent triangular face on the FreeSurfer "white" surface (refined gray/white matter boundary estimate). That single-vertex sum is not exactly constant across vertices because of slight non-uniformities in the final relaxed state of the surface tessellation. However, the sum of vertex-wise areas over a connected region of vertices exactly represents the summed original area of the enclosed triangles (plus the 1/3 fraction of triangles associated with the boundary vertices; along a straight edge of vertices, this last contribution corresponds to half of the area of the triangles just beyond the edge). The minimum areal increment that can be measured is roughly the average original vertex-wise area, which is ~0.6 sq mm.

## Results

We first discuss the amplitude of the vertex-wise response for all relevant Scene Comprehension contrasts. The next set of results present the overlap of scene activation with retinotopic, tonotopic, and somatomotor maps. Finally we discuss scene activation relative to reading. The main scene comprehension contrast utilized for overlap analysis with sensory-motor maps is Story vs. Jigsawed contrast. The scene comprehension activation is illustrated as transparent overlays over single modality phase hue maps. For clarity, in later overlap figures, only positive activation (for Story vs. Jigsawed) after thresholding and cluster correction is shown. The overlap results for different modalities are illustrated for three individual subjects and then for the group as a whole. For the cross-subject average, the sensory-motor maps are illustrated for two separate vertex thresholds; p<0.05 (lower threshold) and p<0.01 (higher threshold), corrected for multiple comparisons using cluster thresholding at p<0.05. The scene comprehension data illustrated here uses a vertex threshold of p<0.01 in all cross-average images. For the individual subjects, results are illustrated at a vertex threshold of p<0.001, corrected to p<0.01 for both scene comprehension and topological mapping data.

For overlap with reading activation, the cross-subject scene comprehension contrast (Story vs. Jigsawed) is illustrated as a transparent overlay over the reading contrast (English vs. Hindi) activation. The scene comprehension and reading group average data are depicted for two vertex thresholds p<0.01 and p<0.05, cluster corrected to p<0.05. A final omnibus figure shows the sensory-motor map locations added as outlines over the scene/language overlap illustration.

All 9 subjects who participated in the scene comprehension experiment gave satisfactory performance in the target detection task, and comprehension assessment, as well as having motion well under our threshold of 1 mm. The activation for the target detection regressor (button press regressor) was used as an extra quality check to decide whether the subject performed the task as per the instructions during each run. All our subjects had comparable performance across conditions in the target detection task and motor activation for the target regressor.

### Target detection response

The target detection response time was calculated as the time it took for participants to respond to the target after the target presentation start time, target was presented for a second after the start time. On average, participants took 1.08 seconds ± 0.01 (SEM) to respond to the target when it occurred in Story, 1.13 seconds ± 0.09 (SEM) in Jigsawed, 1.08 seconds ± 0.05 (SEM) in Shuffled and 1.38 ± 0.06 (SEM) in Fixation. A Wilcoxon matched-pairs signed-ranks test indicated no significant differences between Story (median= 1.06 seconds) and Jigsaw (median=1.08 seconds) (Z=-0.06, p=0.95) or between Story and Shuffled (median = 1.31 seconds; Z=-0.42, p=0.68). The differences between Story and Fixation (median = 1.31 seconds) was found to be significant (Z=-2.67, p=0.008).

On average, participants managed to detect 73% ± 0.06 (SEM) targets in the Story condition, 80.9% ± 0.03 (SEM) in Jigsawed condition, 79.6% ± 0.07 (SEM) in Shuffled condition and 93.6% ± 0.03 (SEM) in Fixation. A Wilcoxon matched-pairs signed-ranks test was carried out to assess statistical significance of the average success rate. The median success rate for targets in Story, Jigsawed, Shuffled, and Fixation were 71.4%, 85.7%, 83.3% and 100% respectively. There were no significant differences between Story and Jigsaw or Story and Shuffled (Story vs. Jigsawed: Z=-1.41, p=0.16; Story vs. Shuffled: Z=-0.771, p=0.44). The performance in Fixation condition was significantly better (Z=-2.2, p=0.03), not surprisingly as the target was easier to discern in the Fixation condition.

### 4.4.2 Scene Comprehension Activation

Figures 2 and 3 illustrate the average cross-subject activation for a stair-step of t-values (all above a minimum threshold of p<0.05, uncorrected) for each condition (Story, Jigsawed and Shuffled) relative to Fixation (Fig. 2), and those for two main contrasts, Story vs. Jigsawed and Story vs. Shuffled (Fig. 3).

**Figure 2:**
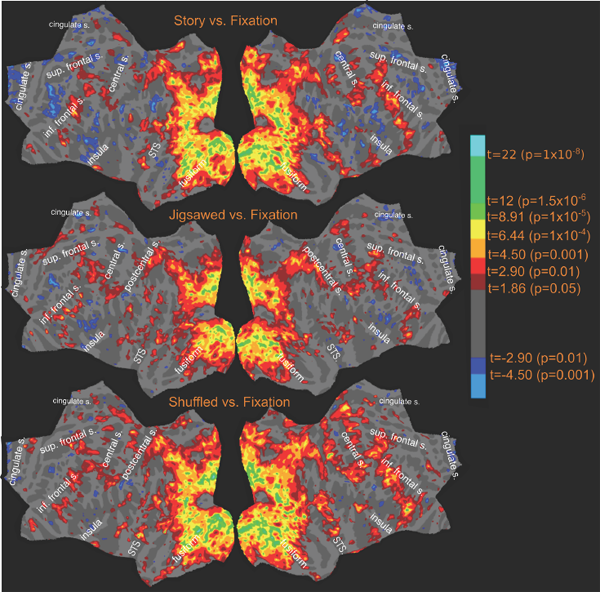
Activation amplitude profile (uncorrected) for relevant contrasts.

**Figure 3:**
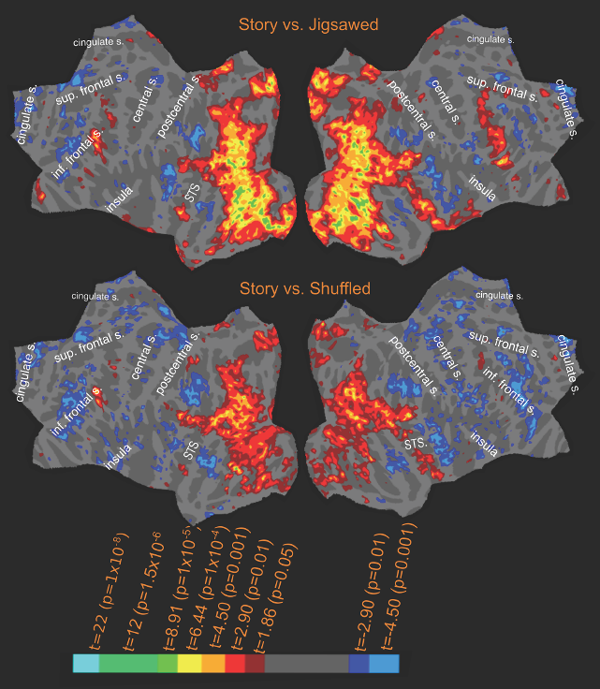
Activation amplitude profile (uncorrected) for relevant contrasts contd.

In the cross-subject activation for Story vs. Jigsawed (Fig. 3, top), the most extensive activation is observed in the posterior occipital cortex. The left hemisphere activation covers the entire lateral occipital cortex. There are several branches which extend anteriorly beyond lateral occipital cortex, one branch extends into the temporal lobe, stretching across MT and posterior STS reaching up to STG. Another offshoot reaches up to the intraparietal sulcus (IPS) covering regions in the superior parietal lobules. The next offshoot stretches nearly half the length of the inferior temporal gyrus and extends ventrally onto the fusiform gyrus joining the medial activation covering most of collateral sulcus and part of parahippocampal cortex, stretching across lingual gyrus, calcarine sulcus and reaching beyond parieto-occipital sulcus. There is also strong activation in the precuneus. Occipital cortex has been known to have several scene selective regions (parahippocampal place area, retrosplenial complex, occipital place area) and object selective regions (lateral occipital cortex) and face selective regions (fusiform face area, occipital face area). The activation observed for Story vs. Jigsawed is consistent with those findings (PPA and RSC locations annotated in figures are based on the functional localizations identified by Nasr et al. (2011) on the fsaverage surface). Regions in occipital cortex which did not show a significant difference in activation between Story and Jigsawed included early visual areas such as V1, V2 (especially the foveal regions) and the adjoining cuneus cortex.

In the left frontal cortex, there are two distinct activation zones — one in the precentral sulcus near FEF (visible at a lower threshold of p<0.05) and another near the inferior frontal sulcus (IFS) in the pars opercularis region. In the left temporal cortex, apart from the occipital branch that extends into lateral temporal cortex across MT, there is a well separated inferior and anterior activation zone in the superior temporal sulcus.

The activation pattern in the right hemisphere, though similar has some notable differences. The posterior occipital cortex activations are similar to their left hemisphere counterparts, but more extensive. In the right temporal cortex, there is significant activation along the entire superior temporal sulcus. The right frontal cortex also exhibits more extensive activation with activation spread along the precentral sulcus.

The single subject activation for Story vs. Jigsawed was very similar to the cross-subject profile.

In the Shuffled condition, subjects saw exactly the same scenes as in the Story condition, but in a different temporal order. This resulted in partial and imperfect comprehension of the story. The activation pattern for Shuffled vs. Fixation is very similar to Story vs. Fixation as expected. Although subtle in the condition vs. OFF depictions in Figure 2, there are significant differences between Story and Shuffled conditions (Figure 3), and the activation pattern for Story vs. Shuffled shows remarkable similarity to the regions activated in Story vs. Jigsawed condition, albeit with a lower significance value. In the left hemisphere, significant activation (Figure 3) was observed in the lateral occipital cortex, in the branch leading up to STG, as well as in the anterior STS and in the inferior frontal cortex. The activations observed in inferior frontal cortex, anterior STS and part of lateral occipital cortex were exclusive to Story and Shuffled conditions and were absent in the Jigsawed condition. All these regions were significantly more active in the Story condition as compared to the Shuffled condition. On the medial wall, the activation profiles of Story and Shuffled (vs. Fixation) look very similar, while Jigsawed had no activation in precuneus and reduced activation in parieto-occipital sulcus and in the region of left PPA. The medial activation for Story vs. Shuffled was more attenuated than the activation in lateral occipital cortex. However, there was less extensive but significant activation in precuneus, and along the parieto-occipital sulcus extending across RSC and the peripheral regions of early visual areas joining up with activation in the vicinity of PPA.

The activation pattern for Story vs. Shuffled in the right hemisphere was similar, with most of lateral occipital and right STS activation still significantly higher for Story than for Shuffled. Much of the activation observed in right frontal cortex for Story vs. Jigsawed was not observed in Story vs. Shuffled condition, and the medial activation was largely reduced, showing the same pattern as in the left hemisphere.

### Overlap of scene (Story vs. Jigsawed) activation with visual, auditory, and somatomotor maps

The main contrast used in the overlap figures to assess scene comprehension was Story vs. Jigsawed. In the figures depicting overlap with sensory-motor maps, transparent bright yellow regions outlined in black depict the regions that showed significantly higher activation when viewing story compared to jigsaw.

For the cross-subject averages, the visual, auditory, and somatomotor activations were assessed for two different hard, vertexwise thresholds before cluster exclusion correction: p<0.01 (higher threshold) and p<0.05 (lower threshold). Scene activation used a single higher hard, vertexwise threshold of p<0.01. For single subject activations, scene comprehension, retinotopic, and tonontopic maps were hard thresholded at p<0.001, corrected to p<0.01. Among the single subjects included below, subject-3 represents the same subject-3 who participated in Reading experiment (Sood & Sereno, 2016). Subjects 1 and 2 are different from the single subjects illustrated in the Reading experiment. The overlap estimates below, are expressed as the percentage of scene comprehension activation intersecting with sensory-motor maps. The results include an overall estimate, where the percentage of total scene comprehension activation overlapping with retinotopic, tonotopic and somatomotor maps is reported for each hemisphere. Additionally, each region (frontal, temporal and occipito-parietal) for scene comprehension activation is considered separately, and corresponding overlap is expressed as a percentage of the regional scene comprehension activation.

#### Scene comprehension overlap with retinotopic maps

The retinotopy/scene comprehension overlap for the cross-subject surface average is shown in Figure 4. Most of the scene comprehension activation in occipito-parietal cortex, falls within retinotopic regions in both hemispheres. Part of the activation in parahippocampal cortex, parieto-occipital sulcus and precuneus on the medial side falls outside the bounds of retinotopy. While the posterior superior temporal activation largely falls within retinotopy, the activation along middle and anterior STS does not overlap with retinotopic maps. There is also significant overlap in the frontal cortex, and the activation near the inferior frontal sulcus partially overlaps with retinotopic maps.

**Figure 4:**
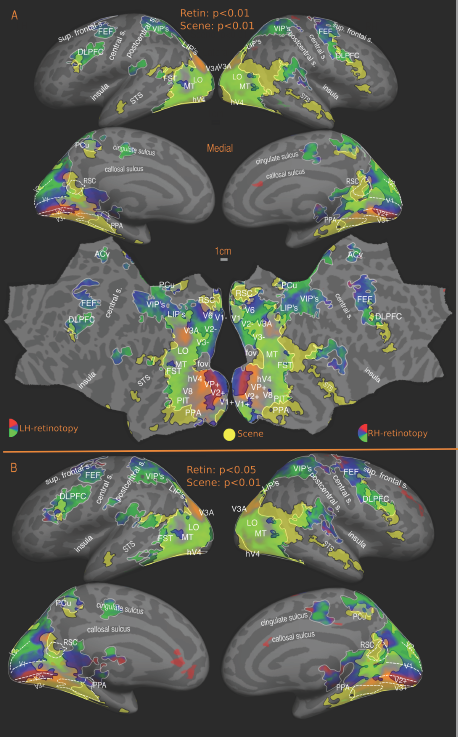
Overlap of scene (Story vs. Jigsawed) with retinotopic maps-group. Reported P values are vertex thresholds used before cluster thresholding. All activations are cluster thresholded to P < 0.05. Scene activation is illustrated at P < 0.01 (vertex threshold) in A and B. Retinotopic activation is illustrated at two different vertex thresholds-P < 0.01 (A) and P < 0.05 (B). In overlap figures (4-8), sensory-motor maps are represented using red, blue and green colors, with map borders outlined in white. Reading activation uses a uniform yellow color with borders indicated using black.

Overall, at a higher threshold, more than 65% of the cross-subject scene comprehension activation in both hemispheres falls within retinotopic areas (LH: 80%, RH: 67%) rising to more than 70% at lower threshold (LH: 85%, RH: 74%). In occipito-parietal cortex, the estimates are 84% in LH and 77% in RH at higher threshold, rising to 89% and 83% when the threshold is lowered. In the temporal cortex, there is only a modest overlap with retinotopy with 3% of LH and 12% of RH activation overlapping with retinotopy at higher threshold rising to 17% and 23% respectively at lower threshold. There is also substantial overlap in the frontal cortex with around 54% of LH and 15% of RH activation falling within retinotopic regions at higher threshold, and rising to 75% and 47% at lower threshold.

For the individual subjects (Figure 5), the overall left hemisphere retinotopy/scene comprehension overlap in LH was 66% for subject-1, 55% for subject-2 and 75% for subject-3, while the right hemisphere overlap was 60% for subject-1, 72% for subject-2 and 56% for subject-3.

**Figure 5:**
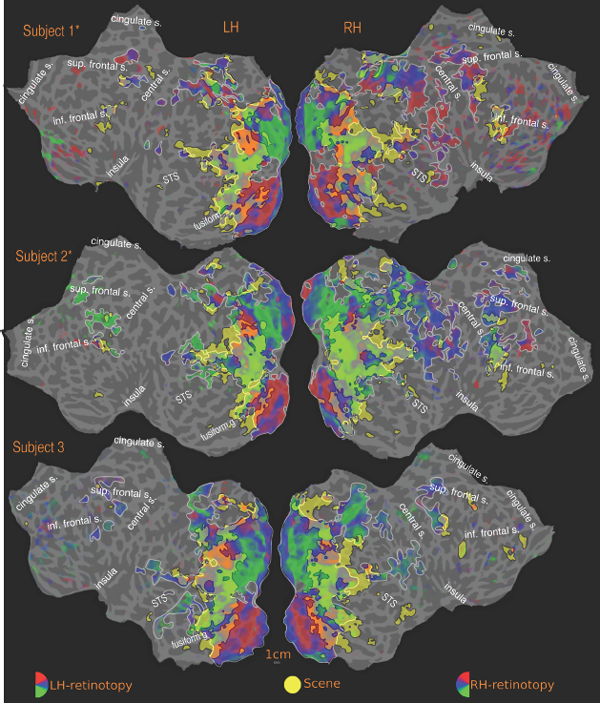
Overlap of scene (Story vs. Jigsawed) with Retinotopic maps-individual subjects. Activations are illustrated at P < 0.001, corrected to P<0.01.

Zooming in on individual regions, as with the cross-subject average, there is more than 60% overlap between scene comprehension and retinotopy in occipito-parietal cortex in both hemispheres (subject-1: 73%, subject-2: 60%, subject-3: 77% in LH and subject-1: 70%, subject-2: 78%, subject-3: 64% in RH). In the left hemisphere frontal and temporal scene activation, overlap with retinotopy is 33% and 13% in subject-1 and 39% and 29% in subject-2 and 18% and 51% for subject-3. The corresponding overlaps for the right hemisphere of these subjects are 27% (frontal) and 33% (temporal) for subject-1, 53% (frontal) and 47% (temporal) for subject-2 and 16% (frontal) and 17% (temporal) for subject-3.

#### Scene overlap with tonotopic maps

In the left temporal cortex, the tonotopic maps partially overlap the activation in posterior STG, but the significant activation in the anterior STS is just outside the bounds of tonotopic maps. The frontal lobe scene comprehension activation also substantially overlaps with frontal tonotopic maps there. Those are also potential sites of multi-sensory integration since the regions contain both retinotopic and tonotopic maps. The individual overlaps are consistent with the cross-subject average overlaps -- that is, there were no idiosyncratic overlaps that disappeared in the average.

The overall tonotopy/scene comprehension overlap (Fig. 6) estimates for cross-subject average are 0.19% (LH) and 1.4% (RH) for the higher threshold, and 1% (LH) and 6% (RH) for the lower threshold in the cross-subject average. In left temporal cortex, the level of overlap is 3% at the higher threshold and 6% at the lower threshold. In the right hemisphere, scene comprehension activation in temporal cortex is more extensive than in the left hemisphere, and the tonotopy overlap estimates are 11% at higher threshold rising to 34% at lower threshold. The tonotopic maps also overlap with frontal scene comprehension activation in both hemispheres: 3% (higher threshold) and 42% (lower threshold) in left hemisphere and 6% (higher threshold) and 60% (lower threshold) in the right hemisphere.

**Figure 6:**
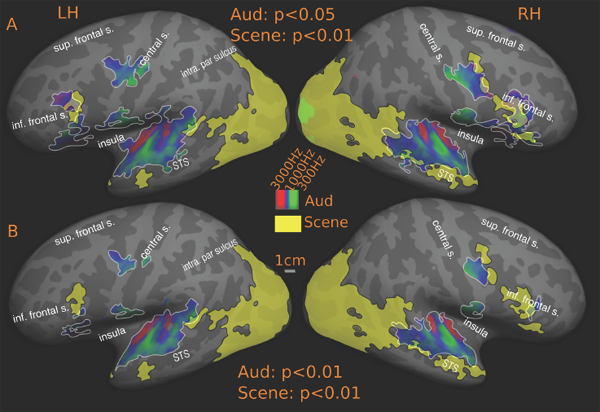
Overlap of scene (Story vs. Jigsawed) with tonotopic maps-Group. Reported P values are vertex thresholds used before cluster thresholding. All activations are cluster thresholded to P < 0.05. Scene activation is illustrated at P < 0.01 (vertex threshold) in A and B. Tonotopic activation is illustrated at two different vertex thresholds-P < 0.05 (A) and P < 0.01 (B).

Among the individual subjects (Fig. 7), subject-1 and subject-2 exhibit a similar profile to the higher threshold group average profile (note that the single subject activations are thresholded at p<0.001), while subject-3 has a much more extensive overlap with tonotopic maps. Overall, around 0.2% in subject-1, 0.5% in subject-2 and 3% in subject-3 overlapped with tonotopy in the left hemisphere. Around 4%, 3% and 9% overlap was observed in the right hemisphere for subject-1, subject-2 and subject-3, respectively. Looking at the regional activations, in subject-1, 2% of left frontal activation and 21% of right frontal activation overlaps with tonotopy. No overlap is observed in left temporal cortex and 18% overlap was observed in right temporal cortex. For subject-2, there is no overlap in the left frontal cortex and a 1% overlap in right frontal cortex, with the corresponding figures in temporal cortex being 5% in left hemisphere and 32% in the right hemisphere. In subject-3, none of the frontal scene comprehension activation in the left hemisphere overlaps with tonotopy, while in the right hemisphere around 40% of frontal activation is within tonotopic maps. The corresponding figures for temporal cortex are 53% in left hemisphere and 56% in the right hemisphere.

**Figure 7:**
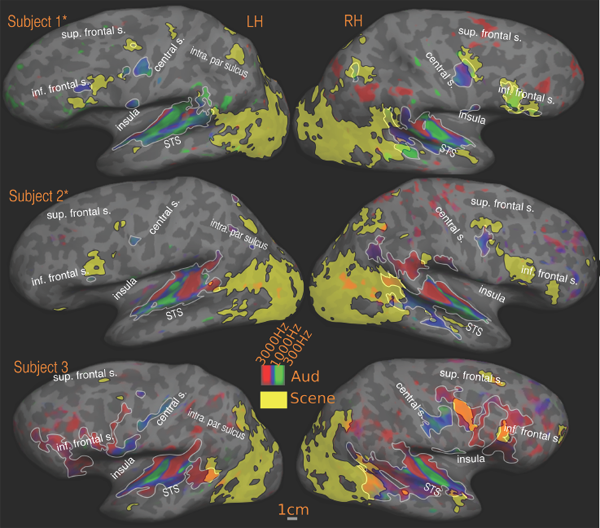
Overlap of scene (Story vs. Jigsawed) with tonotopic maps-individual subjects. Activations are illustrated at P<0.001, corrected to P<0.01.

### Scene comprehension overlap with somatomotor maps

In the cross-subject maps (Fig. 8), there is no overlap between scene comprehension and somatomotor activation in the left hemisphere. The precentral scene comprehension activation (at p<0.05, Fig. 3) lies just outside the region representing the mouth in the somatomotor maps. In the right hemisphere, the precentral region overlaps the outer edge of that region.

**Figure 8:**
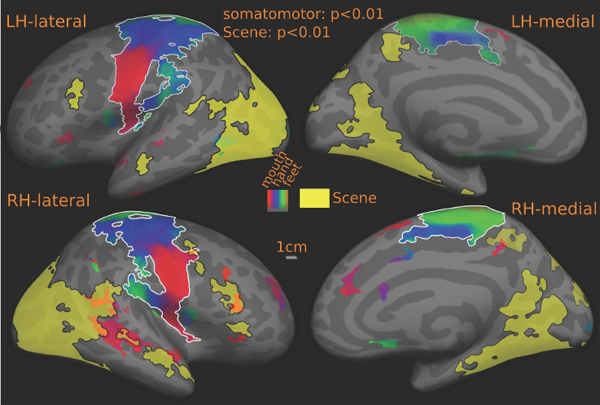
Overlap of scene (Story vs. Jigsawed) with somatomotor maps-Group. Reported P values are vertex thresholds used before cluster thresholding. All activations are illustrated at a vertex threshold of P< 0.01 and cluster thresholded to P < 0.05.

### Comparison between Scene (Story vs. Jigsawed) and Reading (English vs. Hindi) comprehension (Figs. 9, 10)

The results support the classical observation that reading is more left lateralized while picture-based comprehension activates right hemisphere regions more extensively. While the extent and spread differs, the activated regions are mostly aligned in both reading and scene comprehension, with the main activity observed in occipital, temporal and frontal cortex. Left-right hemisphere differences are more prominent with left-lateralized Reading than with right-lateralized scene comprehension.

In occipito-parietal cortex, the common activated regions include V8, hV4, MT, posterior IT, fusiform gyrus, peripheral V1 and V2 and the LIP (lateral intraparietal) regions. All the activated regions in occipito-parietal cortex common to the reading and scene comprehension contrasts fall within retinotopic maps. Unique reading activation in the occipital cortex is mainly in the foveal regions in the primary visual areas, while occipital cortex scene activation is more peripheral. All reading-only activation in the occipital cortex falls within retinotopy. There are scene comprehension-only activations also in occipital cortex which are not shared with reading. In the left hemisphere, where reading activation is more extensive, the regions unique for scene comprehension include activations in parahippocampal cortex (PPA region), RSC, precuneus and in LO. Among these, PPA, RSC and precuneus activations are partially covered by retinotopy. The scene comprehension-only LO region and the surrounding lateral regions shared with reading is also active in the Story vs. Shuffled contrast. In the right hemisphere, there is very little lateral occipital activation for reading, while scene comprehension activation there is merely more spread out than its left hemisphere counterpart. Apart from the regions mentioned in left hemisphere, the scene comprehension regions that do not overlap with retinotopy include regions beyond MT in the superior posterior MTG area.

In the temporal cortex, reading showed prominent activation in the left hemisphere, with extensive activation along the superior temporal gyrus/sulcus and the middle temporal gyrus. The main overlap between reading and scene comprehension is in the posterior superior STS/STG. The more anterior/inferior activation zone in left STS for scene comprehension is largely non-overlapping with reading activation. In the right hemisphere, temporal activation is more extensive for scene comprehension, and although there is a good degree of overlap with reading activation in the STS, there is considerable activation in the anterior STS that is outside the bounds of reading activation.

The frontal activation zones for both reading and scene comprehension are well aligned. The frontal activation in left hemisphere is much more extensive for reading than for scene. There is a prominent overlap near inferior frontal sulcus in the pars opercularis region, as well as an overlap at a less significant (p<0.05) threshold in the precentral sulcus near the FEF region, slightly anterior and inferior to PZ (Huang and Sereno, 2007). In the right hemisphere, scene comprehension activation (Story vs. Jigsawed) is more extensive and is spread along the precentral sulcus all the way up to the pars triangularis region.

The medial cingulate region activated for reading (only present when reading English) and overlapping with a tonotopic map in the region (Figures 9, 10), is not activated for any conditions in the scene comprehension experiment. This could be a bonafide language-specific region.

**Figure 9:**
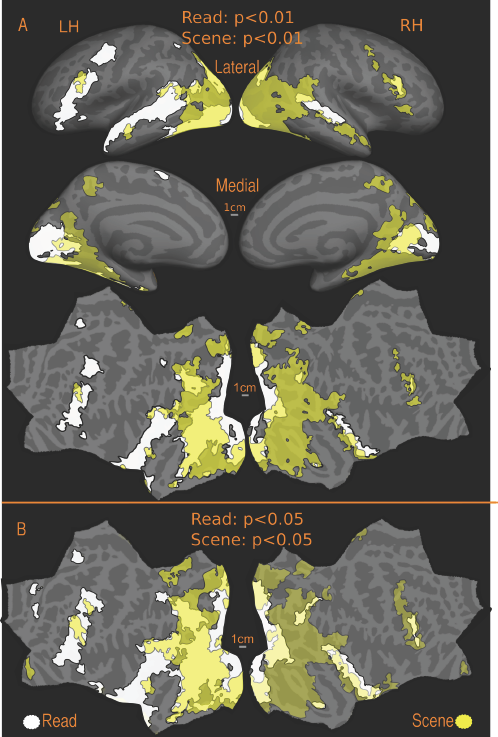
Overlap of scene (Story vs. Jigsawed) with reading (English vs. Hindi) activation-Group. Reported P values are vertex thresholds used before cluster thresholding. All activations are cluster thresholded to P < 0.05. Scene and Reading activations are illustrated at P < 0.01 (vertex threshold) in A and at P<0.05 in B.

**Figure 10:**
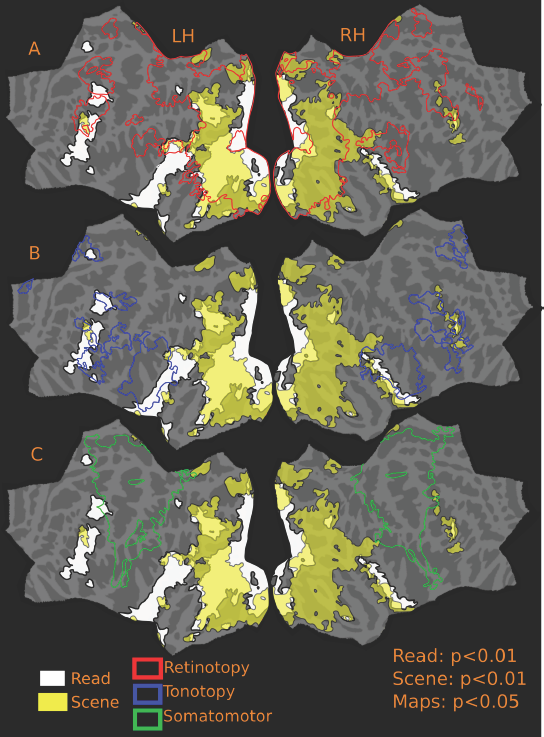
Overlap of scene (Story vs. Jigsawed) and reading (English vs. Hindi) with all maps-Group. Scene and reading activation and are illustrated at vertex threshold P < 0.01, while maps are illustrated at vertex threshold of P<0.05. All activations are cluster corrected to P < 0.05

## Discussion

The study presented in this paper compares and contrasts naturalistic narrative scene and reading comprehension with topological visual, auditory and somatomotor maps. There were two main objectives. The first was to accurately localize regions of interest in narrative scene comprehension by determining their exact relation to low and high-level topological visual, auditory, somatosensory and motor maps. The figures presented here are the first illustrations of the relative location of ‘narrative scene comprehension’ activation and topological visual, auditory and somatomotor maps across the entire cortex in the same group of subjects. Additionally, the results provide a quantitative estimate for the level of overlap between activations observed during narrative scene comprehension and topological sensory-motor maps, which can be driven and outlined by relatively low-level sensory-motor stimuli. At a higher, more rigorous threshold of p<0.01, nearly 80% of cross-subject scene activations in left hemisphere and 67% in right hemisphere fall within regions containing topological sensory-motor maps. When the threshold for sensory-motor maps is lowered (p<0.05), these figures rise to 85% and 74% in left and right hemispheres respectively. The second main objective was to compare and contrast the activation during narrative scene comprehension with that observed during narrative reading comprehension. The reading and scene comprehension stimuli both used exactly controlled saccade sequences and naturalistic serial comprehension of story content, making their joint analysis -- together with topological cortical maps in all main modalities -- the first such available complete data set.

The scene comprehension experiment described in this paper differs from other scene comprehension/natural movie studies in the literature in several respects. First, the study’s main focus was comprehension of coherent narrative picture stories unfolding over 30 to 50 seconds, and subjects were tested for their comprehension afterwards. Second, in contrast to majority of neuroimaging studies that use rapid serial visual presentation, the presentation mode used here allowed naturalistic eye movements that were nevertheless carefully controlled across conditions. Finally, the analysis method utilized cortical surface-based group averaging, which introduces less blurring, rather than volume based group analysis commonly employed in majority of scene comprehension studies.

### Activation patterns in Frontal cortex

The frontal activations for both reading and scene comprehension are mostly aligned, but with several unique features. In the left hemisphere, reading activation is more extensive than scene comprehension activation and fully contains the scene comprehension activation there, while in the right hemisphere the reverse is true. In the left hemisphere, the shared activation is found near the inferior frontal sulcus and is mostly contained within the retinotopic and tonotopic maps. This common activation zone also appeared in the Story vs. Shuffled condition. Considering that this region was active for reading (English vs. Hindi), scene comprehension (Story vs. Jigsawed), and in the Story vs. Shuffled contrasts, this could be a candidate region relevant for serial narrative comprehension common to all of these three contrasts. Turning to the right frontal cortex, reading activation is a subset of scene activation; and again the shared activation is contained mainly within tonotopic maps.

Another shared region in the left lateral frontal cortex, PZ, is situated just lateral to the FEF, where both reading and scene activations (p<0.05) are present along with topological maps in all modalities. This is the only region where all three modality maps overlap (there are several other regions where maps from two modalities overlap).

Finally, the dorsomedial frontal eye fields in the medial cortex is yet a region activated for all conditions (both experimental and control) in both experiments (reading and scene comprehension), and as already discussed, this region overlaps with retinotopy.

Fronto-parietal regions are often considered multi-domain sites associated with cognitive processes that are shared across domains (e.g. attention, see Duncan et al., 2000). There is considerable evidence in the literature for frontal activation during many kinds of language tasks. But much less work has been done to identify the relevance of these regions for picture based comprehension. Studies using full movies and silent movies have reported activation in frontal cortex, although where these activations fall relative to language activations or to the attentional network has not been well defined, especially within the same subjects.

The results presented here show that not all activation in frontal cortex during reading is shared with scene comprehension activation and vice versa. The reading (English vs. Hindi) and scene (Story vs. Jigsawed) contrasts used here were controlled for eye movements and were designed such that their attentional demands were similar. Hence the distinct reading/scene regions identified here are candidate regions for specialized linguistic processing or scene processing. Our data suggest that left dorsolateral frontal cortex exhibits more extensive activation during reading that is not present during scene processing, even when the sequential scene processing required similar attentional and working memory demands. Similarly, in the right frontal cortex, scene comprehension activation supersedes activation during reading. Finally, there is also a distinct region in the left middle-anterior cingulate cortex for reading, which overlaps with a tonotopic map found in the region. This activation only appears while reading English and is absent for all other conditions in the reading stimuli as well as in all conditions in scene experiment. This could be a bonafide language-specific area, which we tentatively named the dorsomedial frontal 'ear' fields (Sood and Sereno, 2016)

Siting these regions relative to topological maps provides a much more precise anatomical localization than those currently available in the literature. Not all activations found in the frontal cortex were distinct, and there were common frontal activation zones in all relevant conditions in the reading experiment as well as in scene comprehension, some of which could be attributed to the participation of the eye control network in multiple forms of serial comprehension as discussed further below.

### Activation patterns in Temporal cortex

There are clear distinctions between activation patterns in the main reading and scene comprehension contrasts in the temporal cortex. In the left temporal lobe, activation is extensive for reading and most of it is not shared with scene comprehension. This distinct continuous activation zone (for reading) covers most of STG and STS. Tonotopic maps overlap this region partially, but a significant portion (50-75%) of this activation does not overlap with any topological map. In the right hemisphere, however, the distinct reading activation observed in STS is completely overlapping with tonotopic maps. There is substantial activation along the posterior bank of STS for scene in the right hemisphere, which is largely non-overlapping with the reading activation and all topological maps. The shared activation between reading and scene is largely limited to the activation that extends across MT into posterior STS/STG cortex. The scene comprehension activation in this region is fully contained within the more extensive reading activation found in the superior temporal cortex. Except for a small region in STS, most of this shared activation in both hemispheres falls within retinotopic and tonotopic maps in the region. The data suggests that left temporal lobe (also well known as the region where classical Wernicke’s area is situated) is more specialized for reading, considering the total lack of scene activation in most of the reading activated regions in the left superior temporal cortex. On the other hand, the right STS region has significant activation along its entire length for scene processing, which falls outside the bounds of any maps.

### Activation patterns in Occipito-Parietal Cortex

There is extensive activation in occipito-parietal regions both for reading and scene comprehension. For reading, as with other parts of the cortex, activation is far more extensive in the left hemisphere. The regions commonly activated for reading and scene comprehension include V8, hV4, MT, posterior IT, fusiform gyrus, peripheral V1 and V2 and LIP (lateral intraparietal) regions. All the shared activations in this region fall within retinotopy. Distinct reading activation is mainly found in the foveal regions in the early visual areas V1, V2 and V3/VP, while scene activation here extends more peripherally, as expected. All distinct reading activation in the occipital cortex falls within retinotopy. Scene comprehension activation is more extensive in the right hemisphere. The regions distinct for scene comprehension include activations in parahippocampal cortex (PPA), RSC, Precuneus, and LO. Confirming previous results in the literature, the activations in PPA, RSC and Precuneus, unique to scene processing are partially covered by retinotopy. The distinct LO region and the surrounding lateral regions shared with reading is also active in the Story vs. Shuffled contrast. Although occipital regions are not often associated with high-level cognition, the fact that significant activation is observed in most of the activated lateral occipital regions for the much closer contrast of Story vs. Shuffled suggest that these regions may be more intimately involved in narrative comprehension than has previously been assumed.

### Activation patterns in the eye control network

All scene and reading conditions included naturalistic eye movements which were carefully matched between the experimental and control conditions. As anticipated, all reading conditions (English, Hindi, Dot) and scene conditions (Story, Jigsawed, Shuffled), exhibit significant activation bilaterally near intraparietal/postcentral sulcus, FEF, and the dorsomedial frontal eye fields -- regions known to be activated by visuospatial attention and eye saccades (Pierrot-Deseilligny et al., 2004; McDowell et al., 2008; Müri, and Nyffeler, 2008; Jamadar et al., 2013; O’Reilly et al., 2013). The activations in the dorsomedial frontal eye fields and intraparietal/postcentral sulcus are fully attenuated in the contrasts-English vs. Hindi and Story vs. Jigsawed. An exception is lateral FEF; while the activation disappears in Hindi vs. Dot contrast (Sood & Sereno, 2016), a good part of it is still highly significant in English vs. Hindi contrast. The lateral FEF activation for Story vs. Jigsawed condition, is much less significant and far less extensive than the activation observed in English vs. Hindi. More work needs to be done to assess what exactly contributes to this differential activation patterns in lateral FEF.

## Conclusion

To summarize, using five separate experiments on same set of subjects analyzed with a fully surface-based analysis stream, we have identified possible candidate regions which are both shared and unique to serial linguistic comprehension and serial non-linguistic visual comprehension, and how they are situated relative to the topological cortical maps in the three main modalities (visual, auditory and somatomotor). Our results suggest that not all activations in frontal cortex can be attributed to domain general processes, and that there are regions within left frontal cortex that are uniquely specialized for reading. Our study also suggests a more substantial role for occipital cortex in high level cognition, as significant activation is observed in lateral occipital regions even for a very close contrast such as Story vs. Shuffled story. The results presented here confirm the significance of superior temporal cortex in language processing, a well known classical finding, and identify the temporal and frontal regions crucially involved in scene processing. Finally, almost all shared activations between scene comprehension and reading, and part of the unique activations for each activity, overlap with maps. Maps at higher levels are known to require attention; nevertheless the fact that the cortex doesn't entirely discard basic topological information until the very highest levels of processing suggests that maps and cognition are not as cleanly separable as most suppose.

## Acknowledgments

We thank Birkbeck/ UCL Neuroimaging centre for support.

